# Prediction of the secundary structure at the tRNASer (UCN) of *Lutzomyia longipalpis* (Diptera: Psychodidae)

**DOI:** 10.1101/261297

**Authors:** Richard Hoyos-Lopez

## Abstract

*Lutzomyia longipalpis* is the main vector of *Leishmania infantum*, the etiological agent of visceral leishmaniasis in America and Colombia. Taxonomically belongs to the subgenus *Lutzomyia*, which includes other vector species that exhibit high morphological similarity to the female species difficult to identify vectors in leishmaniasis foci and suggesting the search for molecular markers that facilitate this task, further researchs with mitochondrial genes, chromosome banding, reproductive isolation and pheromones evidence the existence of species complex. The aim of this study was to predict the secondary structure of mitochondrial transfer RNA serine (tRNASer) for UCN codon of *Lutzomyia longipalpis* as molecular marker for identify of this species. Sequences recorded in Genbank of *L. longipalpis* sequences were aligned with tRNA′s from previously described species and then tRNASer secondary structure was inferred by software tRNAscan-SE 1.21. The length of tRNASer was 67 base pairs (bp). Two haplotypes were detected in the five sequences analyzed. The *L. longipalpis* tRNASer showed 7 intrachain pairing in the acceptor arm, 3 in the DHU arm, 4 in the anticodon arm and 5 in the TψC. The size of the loops corresponded to 5 nucleotides in the DHU, 7 in the anticodon, 4 in the variable and 7 in the TψC. *L. longipalpis* is distinguished from other species at subgenera *Lutzomyia* by the secondary structure and substitutions inferred tRNASer evidenced in the primary sequence.

---

*Lutzomyia longipalpis* (Lutz & Neiva, 1912), is a neotropical phlebotomine with a wide and discontinuous geographical distribution linked to areas of tropical dry forest, from southern Mexico to northern Argentina (Arrivillaga 2002, Lainson and Rangel 2005), and It is considered the main vector of *Leishmania infantum* (Nicolle, 1908), a pathogen responsible for the clinical manifestations associated with visceral leishmaniasis in America (Grimaldi 1989, Soares and Turco 2003).

In Colombia, it is responsible for the active transmission of *L. infantum* in epidemic outbreaks of visceral leishmaniosis in Santander, Cundinamarca, Córdoba and Huila (Morrison et al 1993, Floréz et al 2006, Fernandez et al 2002, Gonzales et al 2006). Studies of experimental infection have proven their susceptibility to infection of viruses belonging to the genera Vesiculovirus, Orbivirus, Flavivirus, Alphavirus, Bunyavirus and Phlebovirus (Jennings and Bormann, 1980, Tesh and Modi, 1983).

Taxonomically *L. longipalpis* belongs to the subgenus *Lutzomyia* (França, 1924), differentiated mainly in the male by morphological characters such as the presence of curved mushrooms in the paramer, a tufo of accessory mushrooms at the base of the gonocoxite and in the female by a spermatheca. Barrel-shaped as long as 4x wide (Young and Duncan 1994), this subgenus includes the majority of *Leishmania* spp. vectors. and presents species whose females are morphologically indistinguishable from *L. longipalpis* (Martins 1978; Vigoder et al 2010), making it difficult to identify them using taxonomic keys if there are no males present in the collection. This species presents morphological variations in abdominal tergos (Mangabeira 1969, Ward et al 1985), originating investigations that have shown a complex of species through studies of reproductive isolation (Lanzaro et al 1993), isoenzymes (Lanzaro et al 1998), chromosomal banding (Yin et al 1999), mitochondrial genes (Soto et al 2001, Arrivillaga, et al 2002) and nuclear genes (Peixoto et al 2001, Lins et al 2002, Lins 2008, Hoyos-Lopez et al 2012), pointing out significant genetic differences at the population level as a consequence of the low flight capacity, little dispersion, and presence of climatic and geographic barriers between the sites where it is found (Morrison et al1993, Arrivillaga et al, 2002), it has been suggested that these factors could be involved in differences in the transmission of L. infantum (Rocha et al, 2011).

In this context, the use of morphological characters associated with spermathecae, the high presence of females, the low number of males in the entomological collections, and the similarity between species of the same subgenus *Lutzomyia*, make difficult the taxonomic identification and vectorial incrimination of species that occur sympatrically in foci of leishmaniasis. An alterative is the search for molecular characters that allow differentiating species morphologically similar to *L. longipalpis* within the subgenus *Lutzomyia*, possible species within the longipalpis complex, and in turn determining the evolutionary relationships between populations (Lanzaro and Warburg 1995).

Recently, the mitochondrial transfer gene for serine (UCN) has been proposed as a molecular marker for the genetic characterization and diagnosis of the species (Vivero et al 2007, Pérez-Doria 2008, Pérez-Doria 2011), being explored in 11 species of phlebotomine with emphasis on species with importance in the transmission of leishmaniosis. In the present article we describe, for the first time, the primary and secondary structure of the mitochondrial transfer RNA for serine that recognizes the codon UCN (tRNASer) of *L. longipalpis* from different localities to differentiate it from other species of the subgenus *Lutzomyia*.

## Materials and methods

To describe the primary and secondary structure of the gene for tRNASer, sequences obtained from specimens of *L. longipalpis* collected by the program of study and control of tropical diseases (PECET), of different geographical origin and registered in Genbank under the access numbers were used: AF403498 (Brazil), AF403497 (Brazil), AF403496 (Costa Rica), AF403495 (Costa Rica), AF403494 (Cundinamarca-Colombia), AF403493 (Cundinamarca - Colombia), AF403492 (Cundinamarca - Colombia). The sequences were aligned with the ClustalW algorithm in Bioedit (Hall1 999) and then analyzed in MEGA 4.1 (Tamura et al 2007) to determine the nucleotide composition and polymorphic sites of the gene. The secondary structure of the tRNASer of *L. longipalpis* was obtained with the program tRNAscan-SE 1.21 (Lowe and Eddy 1997) and was manually graphed. Additionally, the sequences of *L. longipalpis* were compared with the sequences described for the genus *Lutzomyia* by Vivero (2007), Pérez-Doria (2008) and Pérez-Doria (2011).

## Results & Discussion

The mitochondrial genome of insects has 13 protein-coding genes, a 12S rRNA gene, a 16S rRNA gene, a control region and 22 tRNA genes (Hoy 2006; Behura 2011), one for each amino acid, with the exception of leucine and serine, which individually possess two types of molecules that recognize the codon AGY and UCN, the latter is the one described in the present study (Hanada et al 2000).

The tRNASer (UCN) of *Lutzomyia longipalpis* has a length of 67 bp, presenting a composition of 43.1% adenine, 7.5% cytosine, 10.7% guanine and 38.8% thymine, highlighting the high proportion of A - T (81.9%) as has been recorded in the mitochondrial genome of arthropods and insects (Crease 1999, Behura 2011). No overlap was observed between the last cytochrome B codon and the first codon of tRNASer, and an intergenic spacer of 8 bp was present between them. This finding contrasts with the economic profile of the mitochondrial genome, the overlap between both genes and the absence of the intergenic spacer in *Lutzomyia trinidadensis* (Newstead 1922), *Lutzomyia dubitans* (Sherlock 1962), *Lutzomyia panamensis* (Shannon 1926) and *Lutzomyia evansi*(Nuñez-Tovar 1924) (Vivero et al 2007, Pérez-Doria et al 2008, Pérez-Doria et al 2011).

The modeled secondary structure of the tRNASer of *L. longipalpis* is made up of the amino acid acceptor arm, the arm and the magnifying glass dihydrouridine (DHU), the arm and the magnifying glass of the anticodon, the variable magnifying glass, the arm and the magnifying glass ribotimidine-pseudouridinacytosine (TψC) (Figure 1). In the acceptor arm of the amino acid, seven matings were observed, six of which corresponded to pair A - U (adenine - uracil) and one to G - C (guanine - cytosine). This acceptor arm of *L. longipalpis* has a similar structure to that inferred for *Lutzomyia hartmanni* (Fairchild and Hertig 1957), highlighting that during the in vivo maturation, the last two bases U - U (uracil - uracil) from the 3 ′end of the tRNASer are replaced by the CCA trinucleotide (cytosine - cytosine - adenine), forming the binding site of the respective amino acid (Pérez-Doria et al 2011).

**Figure 1.**
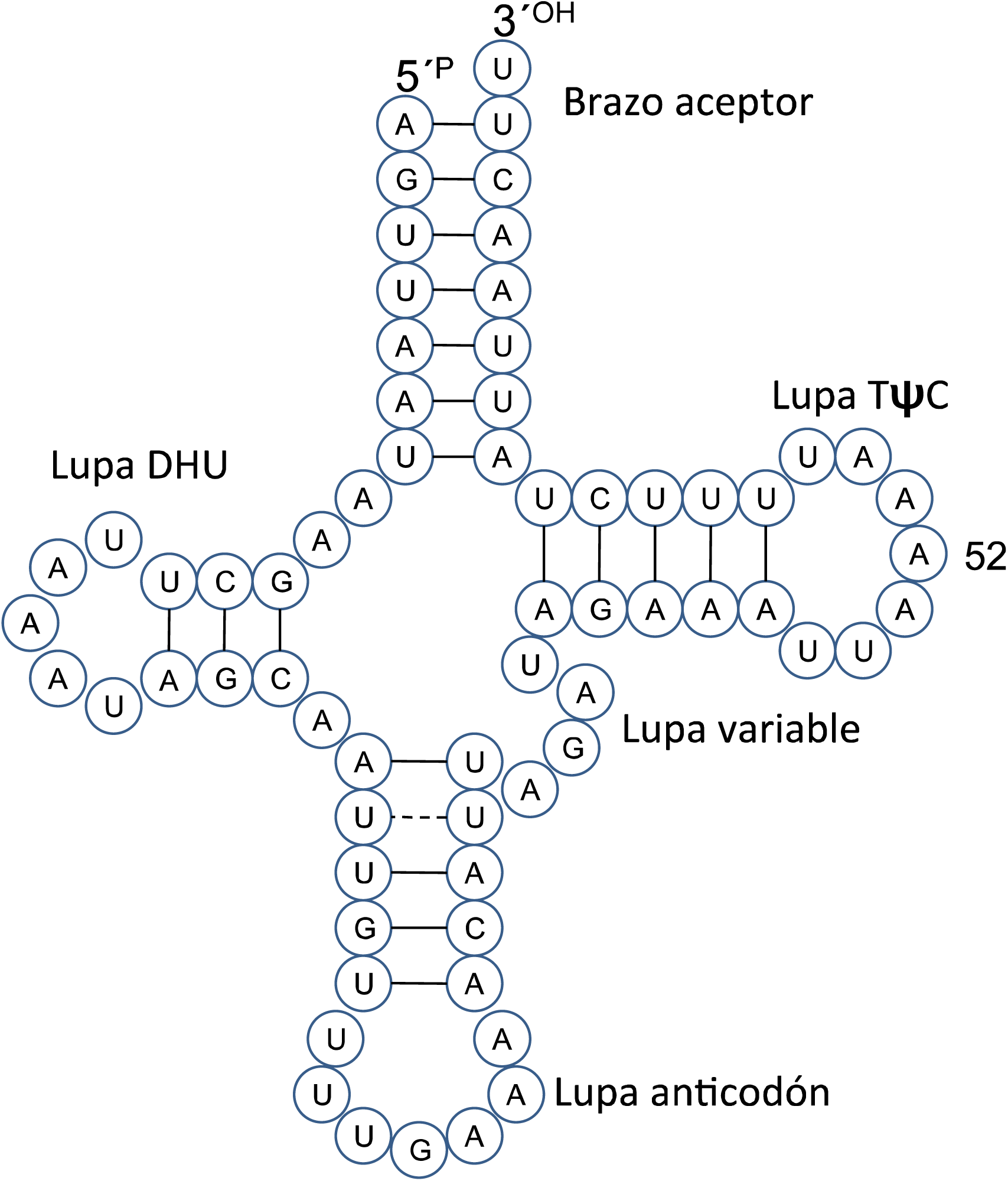
Secondary structure of mitochondrial transfer RNA for serine (codon UCN) of *Lutzomyia longipalpis*. (-), canonical matings. (-) non-canonical matings.

**Figure 2.**
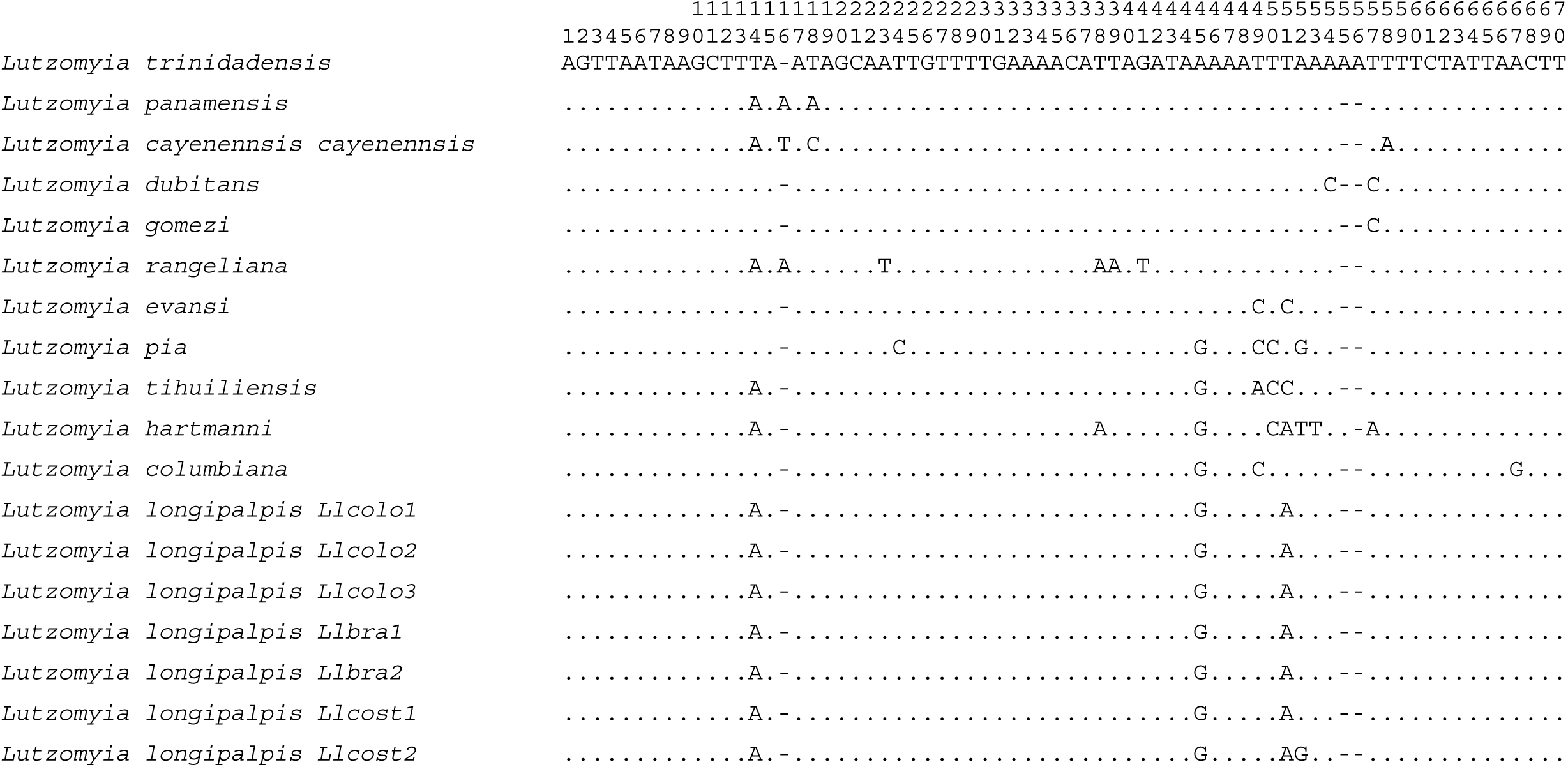
Multiple alignment of the sequences registered in Vivero (2007), Pérez-Doria (2008) and Pérez-Doria (2011). The nucleotides are represented by the first letter of the respective name. The points indicate homology and the dashes correspond to insertion-deletion events.

The number of matings in the DHU arm was three: two pairs G - C (guanine - cytosine) and one U - A (uracil - adenine), and the T□C arm presented five matings, of which four corresponded to pair U - A (uracil - adenine) and a pair G - C (guanine - cytosine). The anticodon arm presented four canonical matings: three corresponding to the pair U - A (uracilo - adenine) and one G - C (guanine - cytosine), the presence of a non - canonical mating U - U (uracil - uracil) was also observed. this arm. The size of the DHU, T□C, variable and anticodon loupes corresponded to five, seven, four and seven nucleotides respectively, also presenting two nucleotides (adenine-adenine) between the acceptor arm and the DHU magnifying glass, and a nucleotide (adenine) between the latter and the anticodon arm.

The tRNASer of *L. longipalpis* is distinguished from the others recorded (Vivero et al 2007, Pérez-Doria 2008, Pérez Doria 2011) mainly by the substitutions in positions 14, 45 and 51; in position 52 a change to guanine is observed, evidenced by a Costa Rican haplotype different to the other analyzed sequences, this change corresponds to adenine to guanine in the TψC magnifying glass and does not originate changes in the secondary structure. By way of conclusion, the primary and secondary structure of the tRNASer of *L. longipalpis* differs from those previously recorded for the genus Lutzomyia, which indicates its usefulness to differentiate it taxonomically, however it is recommended to extend the use of this marker to species that are evolutionarily close to *L. longipalpis* within the subgenus Lutzomyia. Exploration of other molecular markers as cytochrome oxidase I – COI is a tool with remarkable results to differentiate mosquito species and sandflies (Hoyos et al 2015a, Hoyos et al 2015b, Toro-Cantillo et al 2018) in rural areas where conditions favor the transmission of Leishmaniasis (Hoyos et al 2013, Hoyos et al 2016, Toro-Cantillo et al 2017)

## Acknowledgments

To the laboratory of biomedical research and molecular biology from Universidad del Sinú.

